# Neuronal hyperexcitability is a DLK-dependent trigger of HSV-1 reactivation that can be induced by IL-1

**DOI:** 10.1101/2020.04.16.044875

**Authors:** Sean R. Cuddy, Austin R. Schinlever, Sara Dochnal, Jon Suzich, Parijat Kundu, Taylor K. Downs, Mina Farah, Bimal Desai, Chris Boutell, Anna R. Cliffe

## Abstract

Herpes Simplex Virus (HSV) establishes a latent infection in neurons and periodically reactivates to cause disease. The neuronal stimuli that trigger HSV reactivation have not been fully elucidated. Here we demonstrate that HSV reactivation can be induced by neuronal hyperexcitability. Neuronal stimulation-induced reactivation was dependent on voltage-gated ion and hyperpolarization-activated cyclic nucleotide-gated (HCN) channels, demonstrating that neuronal activity is required for reactivation. Hyperexcitability-induced reactivation was dependent on the neuronal pathway of DLK/JNK activation and progressed via an initial wave of viral gene expression that was independent of histone demethylase activity and linked to histone phosphorylation. IL-1β induces neuronal hyperexcitability and is released under conditions of stress and fever; both known triggers of clinical HSV reactivation. IL-1β induced histone phosphorylation in sympathetic neurons, and importantly HSV reactivation, which was dependent on DLK and neuronal excitability. Thus, HSV co-opts an innate immune pathway resulting from IL-1 stimulation of neurons to induce reactivation.

## Introduction

Herpes simplex virus-1 (HSV-1) is a ubiquitous human pathogen that is present in approximately 40-90% of the population worldwide^1^. HSV-1 persists for life in the form of a latent infection in neurons, with intermittent episodes of reactivation. Reactivation from a latent infection and subsequent replication of the virus can cause substantial disease including oral and genital ulcers, herpes keratitis, and encephalitis. In addition, multiple studies have linked persistent HSV-1 infection to the progression of Alzheimer’s disease^2^. Stimuli in humans that are linked with clinical HSV-1 reactivation include exposure to UV light, psychological stress, fever, and changes in hormone levels^3^. How these triggers result in reactivation of latent HSV-1 infection is not fully understood.

During a latent infection of neurons, there is evidence that the viral genome is assembled into a nucleosomal structure by associating with cellular histone proteins^4^. The viral lytic promoters have modifications that are characteristic of silent heterochromatin (histone H3 di- and tri-methyl lysine 9; H3K9me2/3, and H3K27me3)^5–8^, which is thought to maintain long-term silencing of the viral lytic transcripts. Hence, for reactivation to occur, viral lytic gene expression is induced from promoters that are assembled into heterochromatin and in the absence of viral proteins, such as VP16, which are important for lytic gene expression upon *de novo* infection. Reactivation is therefore dependent on the host proteins and the activation of cellular signaling pathways^3^. However, the full nature of the stimuli that can act on neurons to trigger reactivation and the mechanisms by which expression of the lytic genes occurs have not been elucidated.

One of the best characterized stimuli of HSV reactivation at the cellular level is nerve-growth factor (NGF) deprivation and subsequent loss of PI3K/AKT activity^9–11^. Previously, we found that activation of the c-Jun N-terminal kinase (JNK) cell stress response via activation of dual leucine zipper kinase (DLK) was required for reactivation in response to loss of NGF signaling. In addition, recent work has identified a role for JNK in HSV reactivation following perturbation of the DNA damage/repair pathways, which also trigger reactivation via inhibition of AKT activity^12^. DLK is a master regulator of the neuronal stress response, and its activation can result in cell death, axon pruning, axon regeneration or axon degeneration depending on the nature of activating trigger^13,14^. Therefore, it appears that HSV has co-opted this neuronal stress pathway of JNK activation by DLK to induce reactivation. One key mechanism by which JNK functions to promote lytic gene expression is via a histone phosphorylation on S10 of histone H3^15^. JNK-dependent histone phosphorylation occurs on histone H3 that maintains K9 methylation and is therefore known as a histone methyl/phospho switch, which permits transcription without the requirement for recruitment of histone demethylases^16,17^. This initial wave of viral lytic gene expression is known as Phase I, and also occurs independently of the lytic transactivator VP16. In addition, late gene expression in Phase I occurs independent of viral genome replication^18,19^. A sub-population of neurons then progress to full reactivation (also known as Phase II), which occurs 48-72h post-stimulus and requires both VP16 and histone demethylase activity^15,20–23^. However, not all models of reactivation appear to go through this bi-phasic progression to reactivation as axotomy results in more rapid viral gene expression and dependence on histone demethylase activity for immediate viral gene expression.

The aim of this study was to determine if we could identify novel triggers of HSV reactivation and determine if they involved a bi-phasic mode of reactivation. We turned our attention stimuli that cause heightened neuronal activity because hyperstimulation of cortical neurons following forskolin treatment or potassium chloride mediated depolarization has previously been found to result in a global histone methyl/phospho switch^24^. Whether this same methyl/phospho switch occurs in different types of neurons, including sympathetic neurons, is not known. Although forskolin has previously been found to induce HSV reactivation,^25–28^, the mechanism by which forskolin induces reactivation is not known. In particular, if it acts via causing increased neuronal activity and/or as a consequence of activation of alterative cAMP-responsive proteins including PKA and CREB. Hyperexcitability of neurons is correlated with changes in cellular gene expression, increased DNA damage^29,30^, and epigenetic changes including H3 phosphorylation^24^. However, DLK-mediated activation of JNK has not been linked to changes in cellular gene expression nor epigenetic changes in response to hyperexcitability. Using a variety of small molecule inhibitors, we found that forskolin induced reactivation was dependent on neuronal activity. In support of a role for neuronal hyperexcitability causing HSV reactivation, independent stimuli known to cause heightened neuronal activity also induced HSV to undergo reactivation. In addition, DLK and JNK activity were required for an initiation wave of viral lytic gene expression, which occurred prior to viral DNA replication and independently of histone demethylase activity, indicating that hyperstimulation-induced reactivation also is bi-phasic

We were also keen to determine whether we could identify a physiological stimulus for HSV reactivation that acts via causing neurons to enter a hyperexcitable state. IL-1β is released under conditions of psychological stress and fever^31–34^; both known triggers of clinical HSV reactivation^35–37^. IL-1β has previously been found to induce heightened neuronal activity^38–40^. However, an intriguing feature of IL-1β signaling is its ability to have differential effects on different cell types. For example, IL-1β is involved in the extrinsic immune response to infection via activation of neutrophils and lymphocytes. In addition, it can act on non-immune cells including fibroblasts to initiate an antiviral response^41,42^, as has previously been described for lytic infection with HSV-1^41^. Given these differential downstream responses to IL-1β signaling, we were particularly interested in the effects of IL-1β treatment of latently-infected neurons. Interestingly, we found that IL-1β was capable of inducing reactivation of HSV from mature sympathetic neurons. Inhibition of voltage-gated sodium and hyperpolarization activated cyclic nucleotide gated (HCN) channels impeded reactivation mediated by both forskolin and IL-1β. Activity of the cell stress protein DLK was also essential for IL-1β-mediated reactivation. We therefore identify IL-1β as a novel trigger from HSV reactivation that acts via neuronal hyperexcitability and highlight the central role of JNK activation by DLK in HSV reactivation.

## Results

### Increased Intracellular Levels of cAMP Induces Reactivation of HSV from Latent Infection in Murine Sympathetic Neurons

Both forskolin and cAMP mimetics are known to induce neuronal hyperexcitation and have previously also been found to trigger HSV reactivation^25–28^. Using a model of HSV latency in mouse sympathetic neurons isolated from the super-cervical ganglia (SCG)^15^ we investigated whether forskolin treatment induced reactivation in this system and the potential mechanism resulting in the initial induction of viral lytic gene expression. Sympathetic SCG neurons were infected with a Us11-GFP tagged HSV-1^43^ at a multiplicity of infection (MOI) of 7.5 PFU/cell in the presence of acyclovir (ACV). After 6 days the ACV was washed out and the neuronal cultures monitored to ensure that no GFP-positive neurons were present. Two days later, reactivation was triggered by addition of forskolin (Figure 1A). As represented in Figure 1B, forskolin can act either extracellularly on ion channels or intracellularly to activate adenylate cyclase^44–46^. Dideoxy-forskolin (dd-forskolin) is a cell impermeable forskolin analog that can act directly on voltage gated ion channels but does not activate adenylate cyclase^44,47^. We found addition of forskolin but not dd-forskolin triggered robust HSV reactivation (Figure 1C). A slight increase in GFP-positive neurons did occur with dd-forskolin treatment compared to mock (approximately 6.5-fold increase compared to a 130-fold increase for forskolin). Based on a Tukey’s multiple comparison test, this change from mock treated neurons was not significant (P=0.07), however, a direct comparison between mock and dd-forskolin using a T-test suggested a significant induction (P=0.03). Therefore, direct stimulation of ion-channels by dd-forskolin may trigger some reactivation. However, maximal reactivation requires forskolin to be enter neurons. In support of increased intracellular levels of cAMP in inducing HSV reactivation, treatment of latently-infected primary neurons with a cAMP mimetic (8-bromo-cAMP) was sufficient to trigger reactivation (Figure 1D). Furthermore, inhibition of adenylate cyclase activity using SQ22, 536^48^ significantly diminished HSV reactivation (Figure 1E). Therefore, activation of adenylate cyclase and subsequent increased intracellular levels of cAMP are required for robust forskolin-mediated reactivation.

**Figure 1.**
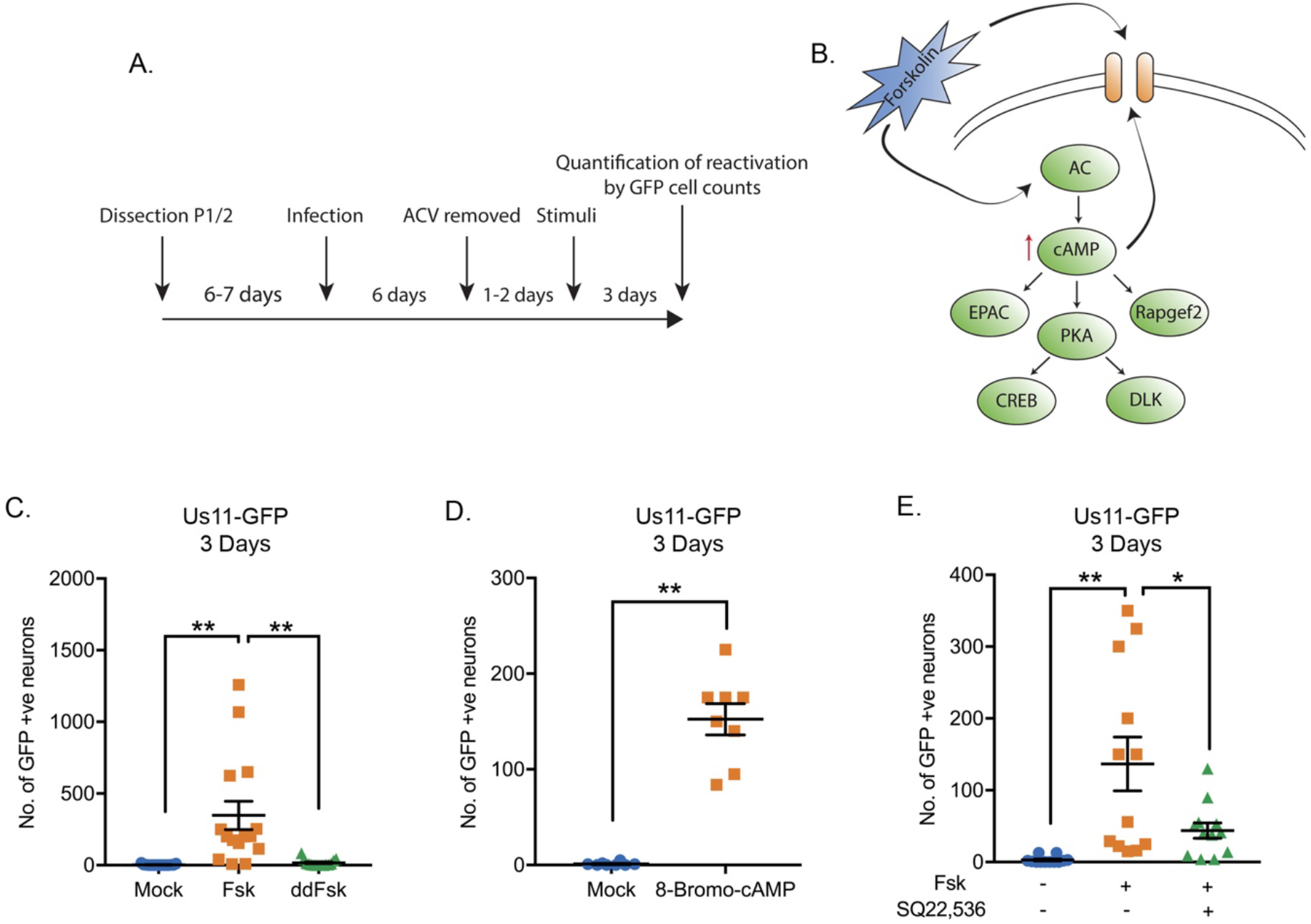
HSV-1 Reactivation Induced by Adenylate Cyclase Activation is DLK/JNK-Dependent. (A) Schematic of the primary superior sympathetic ganglia-derived model of HSV latency. Reactivation was quantified based on Us11-GFP positive neurons in presence of WAY-150168, which prevents cell-to-cell spread. (B) Schematic of the cellular pathways activated by forskolin treatment. Forskolin can act both intracellularly to activate adenylate cyclase increasing the levels of cAMP or extracellularly on ion channels. (C) HSV reactivation was induced by treatment with forskolin (60μM) but not the cell-impermeable dideoxy-forskolin (60μM). (D) Reactivation could be triggered by the cAMP mimetic 8-Bromo-cAMP (125μM). (E) Forskolin-induced reactivation was blocked by the adenylate cyclase inhibitor SQ22,536 (50μM). Each point represents a single replicate. In D statistical comparisons were made using an unpaired t-test. In C and E statistical comparisons were made using a one-way ANOVA with a Tukey’s multiple comparisons test. *<0.05, **P<0.01.

### DLK and JNK Activity are Required for the Early Phase of Viral Gene Expression in Response to Forskolin Treatment

We previously found that DLK-mediated JNK activation was essential for Phase I reactivation following interruption of nerve growth factor signaling^15^. To determine whether DLK and JNK activation were crucial for reactivation in response to hyperexcitability, neurons were reactivated with forskolin in the presence of the JNK-inhibitor SP600125 (Fig. 2A) or the DLK inhibitor GNE-3511^49^ (Fig. 2B). Both the JNK- and DLK-inhibitors prevented forskolin-mediated reactivation based on the number of GFP-positive neurons at 3-days post-stimulus. These data therefore indicate hyperexcitability-induced reactivation is dependent on the neuronal stress pathway mediated by DLK activation of JNK.

**Figure 2.**
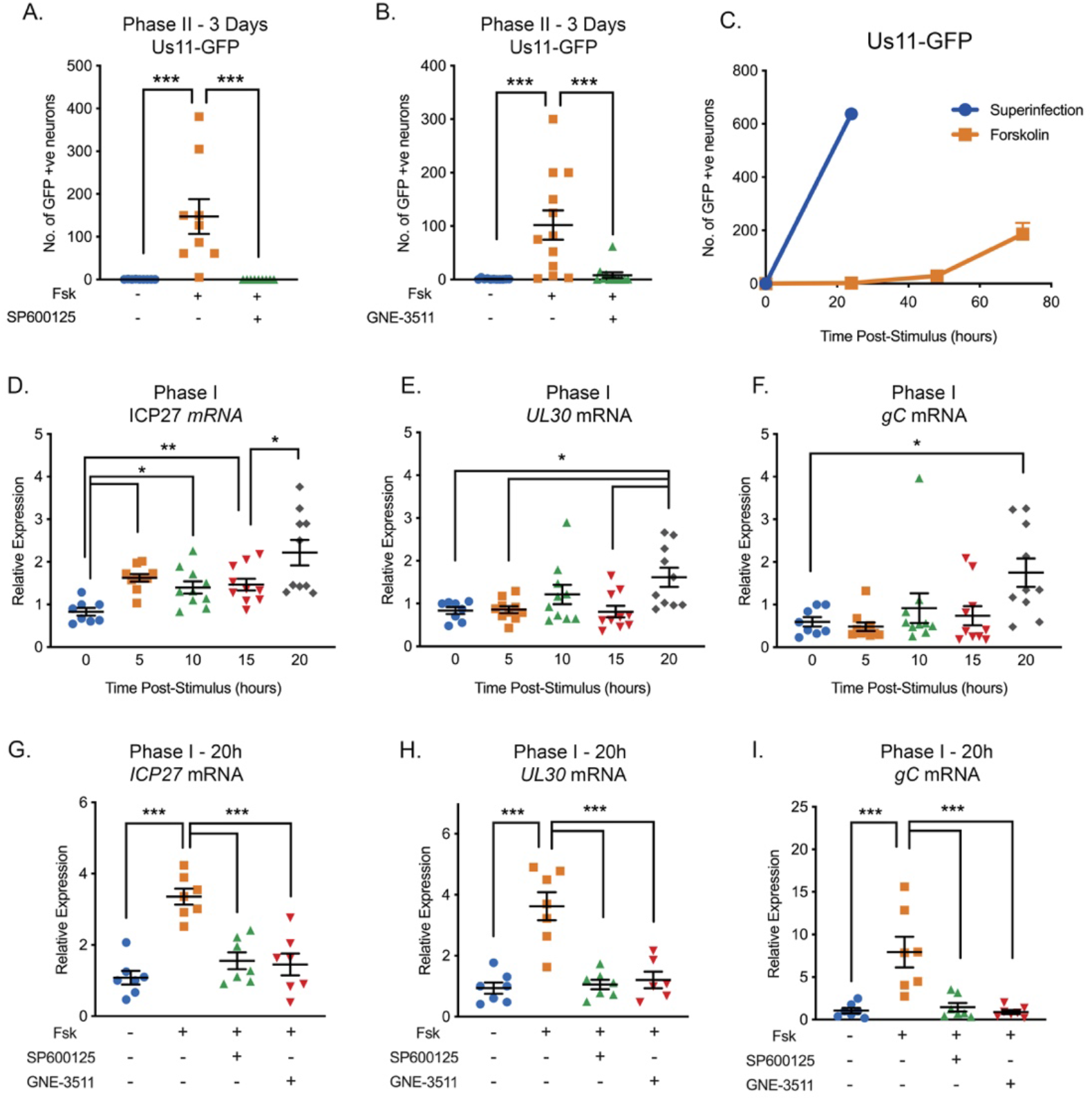
Reactivation Triggered by Forskolin Involves a DLK/JNK Dependent Phase I of Viral Gene Expression. (A) Reactivation was induced by forskolin in the presence of JNK inhibitor SP600125 (20μM). (B) Reactivation was induced by forskolin in the presence of the DLK inhibitor GNE-3511 (4μM). (C) Reactivation was induced by forskolin or superinfection with a wild-type (F strain) HSV-1 (MOI of 10 PFU/cell) and qualified based on Us11-GFP positive neurons (n=3). (D-F) RT-qPCR for viral mRNA transcripts following forskolin treatment of latently infected SCGs. (G-I) RT-qPCR for viral lytic transcripts at 20h post-forskolin treatment and then in presence of the JNK inhibitor SP600125 (20μM) and the DLK inhibitor GNE-3511 (4μM). In D-L each experimental replicative is represented. Statistical comparisons were made using a one-way ANOVA with a Tukey’s multiple comparison. *P<0.05, ** P<0.01, ***P<0.001.

Because we previously found that JNK-activation results in a unique wave of viral gene expression in response to inhibition of nerve-growth factor signaling, we were especially intrigued to determine whether hyperexcitability triggers a similar wave of JNK-dependent viral gene expression. The previously described bi-phasic progression to viral reactivation is characterized by viral DNA replication and production of infectious virus, occurring around 48-72h post-stimulus^18^, but with an earlier wave of lytic gene expression occurring around 20h post-stimulus. To determine whether forskolin-mediated reactivation results in a similar kinetics of reactivation, we investigated the timing of Us11-GFP synthesis, viral DNA replication, production of infectious virus, and lytic gene induction following forskolin treatment. In response to forskolin treatment, Us11-GFP synthesis in neurons started to appear around 48h post-reactivation, with more robust reactivation observed at 72h (Figure 2C). In contrast to forskolin-mediated reactivation, the number of GFP-positive neurons following superinfection with a replication competent wild-type virus resulted in a rapid induction of GFP-positive neurons by 24h post-superinfection (Figure 2C). Therefore, forskolin-triggered reactivation results in slower synthesis of Us11-GFP than superinfection. In addition, these data highlight the ability of forskolin to trigger reactivation from only a subpopulation of latently infected neurons (approximately 1 in every 3.4 neurons compared to superinfection).

The production of infectious virus also mirrored the data for the detection of Us11-GFP positive neurons, with a robust increase in viral titers between 24 and 60h post-stimulus (Figure S1A). An increase in viral genome copy number was also not detected until 48h post-stimulus, which continued between 48h and 72h (Figure S1B). The quantification of viral genome copy number was also carried out in presence of WAY-150138^50^, which prevents packaging of the viral genome^51^, therefore indicating that DNA replication occurs in reactivating neurons and not as a consequence of cell-to-cell spread.

Given the observed 48h delay in viral DNA replication and production of infectious virus, we were interested to determine if there was a Phase I wave of lytic gene expression that occurred prior to viral DNA replication. We therefore carried out RT-qPCR to detect representative immediate-early (*ICP27* and *ICP4*), early (*ICP8* and *UL30*), and late (*UL48* and *gC*) transcripts between 5- and 20-hours post addition of forskolin (Figures 2D-F and S1C-E). For all six transcripts, a significant up-regulation of mRNA occurred at 20h post-treatment, including the true late gene *gC*, whose expression would usually only be stimulated following viral genome replication in the context of *de novo* lytic replication. Therefore, this indicates that lytic gene expression is induced prior to viral DNA replication and that neuronal hyperexcitability does trigger a Phase I wave of lytic gene expression. Notably, we did detect small but reproducible induction of *ICP27* mRNA at 5h post-stimulus, followed by a second induction at 20h (Figure 2D), indicating that there is likely differential regulation of some viral lytic transcripts during Phase I reactivation induced by hyperexcitability that is distinct from both NGF-deprivation and *de novo* lytic infection.

To determine whether JNK and DLK were required Phase I gene expression in response to hyperexcitability, we investigated viral mRNA levels following forskolin-mediated reactivation in the presence of the JNK inhibitor SP600125. We found a significant reduction in *ICP27* (2.2-fold), *UL30* (3.3-fold) and *gC* (5.5-fold) mRNA levels at 20h post-stimulus in the presence of SP600125 (Figure 2G-I). For all genes tested, there was no significant increase in mRNAs in the JNK-inhibitor treated neurons compared to mock. We observed comparable results following treatment with the DLK inhibitor GNE-3511, with a 2.3-, 3-, 8.8-fold decrease in *ICP27*, *UL30* and *gC* mRNAs respectively compared to forskolin treatment alone, and no significant increase in mRNA levels compared to the reactivated samples (Figure 2G-I).

It is possible that in addition to JNK, other signal transduction proteins are important in forskolin-mediated reactivation. Previous data has found that DLK can be activated by PKA, which is well known to be activated by cAMP^52^. However, using well characterized inhibitors of PKA, along with the PKA-activated transcription factors CREB, in addition to two other cAMP responsive proteins Rapgef2 and EPAC, we did not find that these cAMP activated proteins were required for Phase I reactivation (Figure S2). Inhibition of PKA or CREB did reduce Phase II reactivation (Figure S2A and C) but had no effect on Phase I (Figure S2B and D, P=0.354 forskolin versus forskolin + KT 5720, P=0.963 forskolin vs. forskolin + 666-15, Tukey’s multiple comparison test). Inhibition of Rapgef2 or EPAC had no effect on HSV reactivation (Figure S2E and F). Taken together, these data suggest that it is hyperexcitability induced by forskolin that induces a Phase I wave of gene expression via activation of DLK and JNK.

### Forskolin Triggers a Phase I Wave of Viral Gene Expression that is Independent of Histone Demethylase Activity

Hyperexcitability results in the propensity of neurons to fire repeated action potentials, and is associated with specific changes in histone posttranslational modifications. The first is physiological DNA damage^29,30^, measured by the intensity of γH2AX staining in neuronal nuclei. Forskolin treatment was associated with an increase in the levels of γH2AX at 5h post-treatment, which resolved by 15h post-treatment (Figure S3A and C), and is therefore indicative of physiological DNA damage and repair, which occurs upon neuronal hyperexcitability. A second reason for probing the DNA damage/repair pathway in response to forskolin treatment is that previously reactivation of HSV from latency has been associated with perturbation of the DNA damage/repair response^12^. In this previous study, both inhibition of repair and exogenous DNA damage resulted in loss of AKT phosphorylation by PHLPP1, which was required for HSV reactivation. Although we did observe increased levels of γH2AX following forskolin treatment, this was not accompanied by a loss of pAKT measured at 15h post-treatment (Figure S3D). This indicates that HSV reactivation in response to forskolin treatment does not involve dephosphorylation of AKT. Therefore, hyperexcitability triggers reactivation via an alternative mechanism that does not feed into AKT phosphorylation.

Previously, we found that Phase I reactivation is accompanied with a JNK-dependent histone methyl/phospho (marked by H3K9me3/pS10) switch on lytic promoters^15^. In cortical neurons, one study has found that hyperexcitability results in increased H3K9me3/pS10^24^. Therefore, we were particularly interested to determine whether forskolin treatment of sympathetic neurons triggered a histone S10 phosphorylation on H3K9me3. Forskolin triggered a transient increase in H3K9me3/S10p at 5h post-treatment that had returned to baseline by 10h (Figure S3A and B). This indicates that, in keeping with cortical neurons, forskolin induces a histone H3K9me3/pS10 methyl/phospho switch on regions on cellular chromatin.

We next sought to determine whether the phospho/methyl switch that arises as a result of hyperexcitability plays a role in Phase I of HSV reactivation. We therefore investigated whether viral genomes were co-localized with H3K9me3/S10p following forskolin treatment. To visualize HSV genomes, viral stocks were grown in the presence of EdC as described previously^53,54^. Click-chemistry was performed on latently infected and neurons following forskolin treatment. As shown in Figure 3A and B, viral genomes co-localized with H3K9me3/pS10 following robust H3K9me3/S10p staining at 5h. The percentage of viral genomes that co-localized with H3K9me3/S10p was significantly increased compared to the mock reactivated samples at 5h and 20h post-forskolin treatment (Figure 3A).

**Figure 3.**
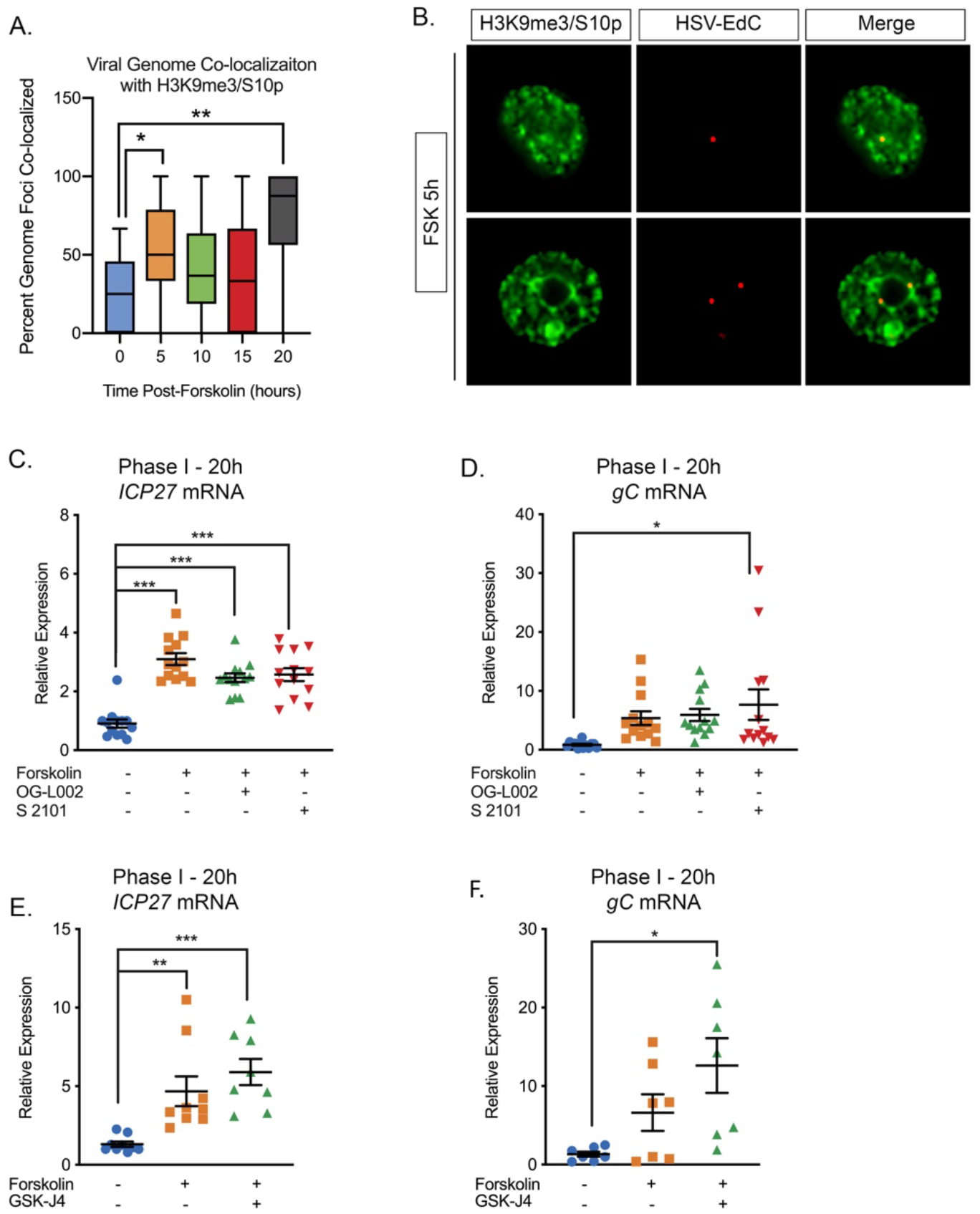
The Initial Wave of Viral Lytic Gene Expression During Forskolin-mediated Reactivation is Independent on Histone Demethylase Activity. (A) Quantification of the percentage of genome foci stained using click-chemistry that co-localize with H3K9me3/S10p. At least 15 fields of view were blindly scored from two independent experiments. Whiskers represent the 2.5-97.5 percentile range. (B) Representative images of click-chemistry based staining of HSV-EdC genomes and H3K9me3/S10p staining at 5h post-forskolin treatment. (C and D). Effect of the LSD1 inhibitors OG-L002 and S 2101 on forskolin-mediated Phase I of reactivation determined by RT-qPCR for *ICP27* (C) and *gC* (D) viral lytic transcripts at 20h post-forskolin treatment and in the presence of 15μM OG-L002 and 20μM S 2102. (E) Effect of the JMJD3 and UTX inhibitor GSK-J4 (2μM) on forskolin-mediated Phase I measured by RT-qPCR for viral lytic transcripts ICP27 (E) and gC (F) at 20h post-forskolin treatment and in the presence of GSK-J4. For C-F each experimental replicate is represented. (C-F). Statistical comparisons were made using a one-way ANOVA with a Tukey’s multiple comparison. *P<0.05, ** P<0.01, ***P<0.001.

Serine phosphorylation adjacent to a repressive lysine modification is thought to permit transcription without the removal of the methyl group^17,24^. Therefore, we investigated whether histone demethylase activity was required for the initial induction in lytic gene expression following forskolin treatment. Previously, the H3K9me2 histone demethylase LSD1 has been found to be required for full HSV reactivation^20,23^, and in our *in vitro* model this was determined by the synthesis of late viral protein at 48-72h post-reactivation^15^. Addition of two independent LSD1 inhibitors (OG-L002 and S 2102) inhibited Us11-GFP synthesis at 72h post-reactivation (Figure S3E). Hence, LSD1 activity, and presumably removal of H3K9-methylation is required for forskolin-mediated reactivation. However, LSD1 inhibition did not prevent the initial induction of *ICP27* and *g*C mRNA expression at 20h post-forskolin treatment (Figure 3C and D). Therefore, this initial wave of viral lytic gene expression following forskolin-mediated reactivation is independent of histone H3K9 demethylase activity.

We previously found that H3K27me demethylase activity is required for full reactivation but not the initial wave of gene expression^15^. However, because of the lack of an antibody that specifically recognizes H3K27me3/S28p and not also H3K9me3/S10p^15^, we are unable at this point to investigate genome co-localization with this combination of modifications. However, we could investigate the role of the H3K27me demethylases in forskolin-mediated reactivation. Treatment of neurons with the UTX/JMJD3 inhibitor GSK-J4^55^ prevented the synthesis of Us11-GFP at 72h post-reactivation, indicating that removal of K27 methylation is required full reactivation (Figure S3F). However, the initial burst of gene expression (assessed by *ICP27* and *gC* mRNA levels) was robustly induced at 20h post-forskolin treatment in the presence of GSK-J4 (Figure 3E and F). Taken together, our data indicate that the initial phase of gene expression following forskolin treatment is independent of histone demethylase activity and therefore consistent with a role for a histone methyl/phospho switch in permitting lytic gene expression.

### Forskolin-Mediated Reactivation Requires Neuronal Excitability

Given that the HSV genome co-localized with regions of hyperexcitability-induced changes in histone phosphorylation, we investigated whether reactivation was linked to neuronal excitability. To inhibit action potential firing, we treated neurons with tetrodotoxin (TTX), which inhibits the majority of the voltage-gated sodium channels and therefore depolarization. Addition of TTX significantly inhibited HSV reactivation triggered by forskolin, as measured by Us11-GFP positive neurons at 72 hours post-stimulus (Figure 4A). To further confirm a role for repeated action potential firing in forskolin-mediated reactivation, we investigated the role of voltage-gated potassium channels, which are required for membrane repolarization. Addition of TEA, which inhibits voltage-gated potassium channel activity, also blocked HSV reactivation measured by Us11-GFP positive neurons at 3 days post-forskolin treatment (Figure 4B). Taken together, these data indicate that action potential firing is required for forskolin-mediated reactivation.

**Figure 4.**
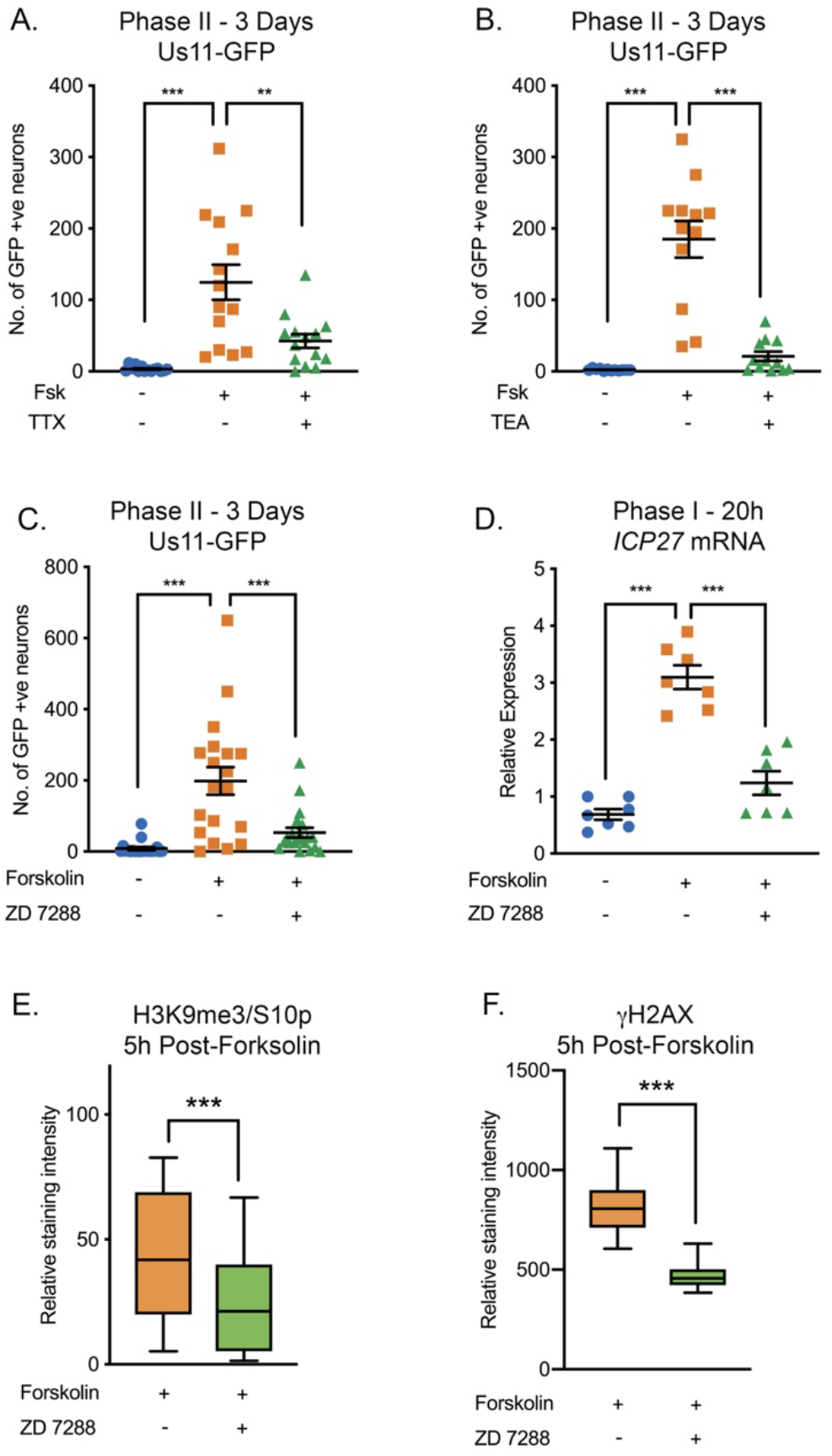
HSV Reactivation Mediated by Forskolin Requires Neuronal Excitability. (A) Latently infected cultures were reactivated with forskolin in the presence of the voltage-gated sodium channel blocker tetrodotoxin (TTX; 1μM) and the number of Us11-GFP positive neurons quantified at 3 days post-reactivation. (B) Latently infected cultures were reactivated with forskolin in the presence of the voltage-gated potassium channel blocker tetraethylammonium (TEA; 10 mM) and the number of Us11-GFP positive neurons quantified at 3 days post-reactivation. (C) Forskolin-mediated reactivation in the presence of the HCN channel blockers ZD 7288 (10μM) quantified as the numbers of Us11-GFP positive neurons at 3 days post-reactivation. (D) The effect of ZD 7288 on the HSV lytic gene transcript ICP27 during Phase I reactivation measured at 20h post-forskolin treatment by RT-qPCR. Individual experimental replicates are represented. (E and F) Quantification of the relative nuclear staining for H3K9me3/S10p and γH2AX in SCG neurons at 5h post-forskolin treatment and in the presence of ZD 7288 from two independent experiments. Statistical comparisons were made using a one-way ANOVA with a Tukey’s multiple comparison (A-D) or two-tailed unpaired t-test (E-F). *P<0.05, ** P<0.01, ***P<0.001.

Increased levels of cAMP can act on nucleotide-gated ion channels, including the hyperpolarization-activated cyclic nucleotide gated (HCN) channels. HCN channels are K^+^ and Na^+^ channels that are activated by membrane hyperpolarization^56,57^. In the presence of high levels of cAMP, the gating potential of HCN channels is shifted in the positive direction, such that HCN channels can open at resting membrane potential, resulting in an increased propensity of neurons to undergo repeated firing^57–59^. HCN channel activity can be blocked by ZD 7288, Ivabradine, or cesium chloride. Addition of ZD 7288 (Figure 4C), Ivabradine (Figure S4A) or CsCl (Figure S4B) all significantly reduced HSV reactivation triggered by forskolin, as measured by Us-11 GFP positive neurons at 3 days post-stimulus. To determine whether HCN channel activity was required for the initial induction of HSV lytic mRNA expression, we assessed viral mRNA expression during Phase I in the presence and absence of ZD 7288. Expression of representative lytic mRNAs *ICP27* (Figure 4D), *UL30* and *gC* (Figure S4C and D) were significantly decreased in the presence of ZD 7288 compared to the forskolin treated neurons alone, and were not significantly increased compared to the mock treated samples. Therefore, HCN channel activity is required for the initial induction of lytic gene expression during Phase I reactivation mediated by forskolin.

We also confirmed that inhibition of HCN-channel activity affected the levels of hyperexcitability-associated changes in histone post-translational modifications. Addition of ZD 7288 resulted in significantly decreased staining intensities of both H3K9me3/S10p and γH2AX at 5h post-forskolin treatment (Figure 4E and 4F), which was the peak time-point for which we observed these changes upon forskolin treatment alone (Figure S3B and S3C). Therefore, activity of the HCN channels in response to increased levels of cAMP, results in hyperexcitability-associated changes in histone modifications and reactivation of HSV from latent infection.

### HSV Reactivation can be Induced by Stimuli that Directly Increase Neuronal Excitability

The role of ion-channel activity in forskolin-mediated reactivation prompted us to investigate whether additional stimuli that induce hyperexcitability in neurons also trigger HSV reactivation. We were also interested in whether reactivation required chronic versus short term hyperexcitability. Increasing the extracellular concentration of KCl is well-known to induce action potential firing. Therefore, we investigate the timing of both KCl and forskolin-mediated hyperexcitability in HSV reactivation. Both of these treatments triggered HSV reactivation more robustly if applied for 8h or more (Figure 5A). This indicates that chronic neuronal hyperexcitability is important in inducing reactivation of HSV.

**Figure 5.**
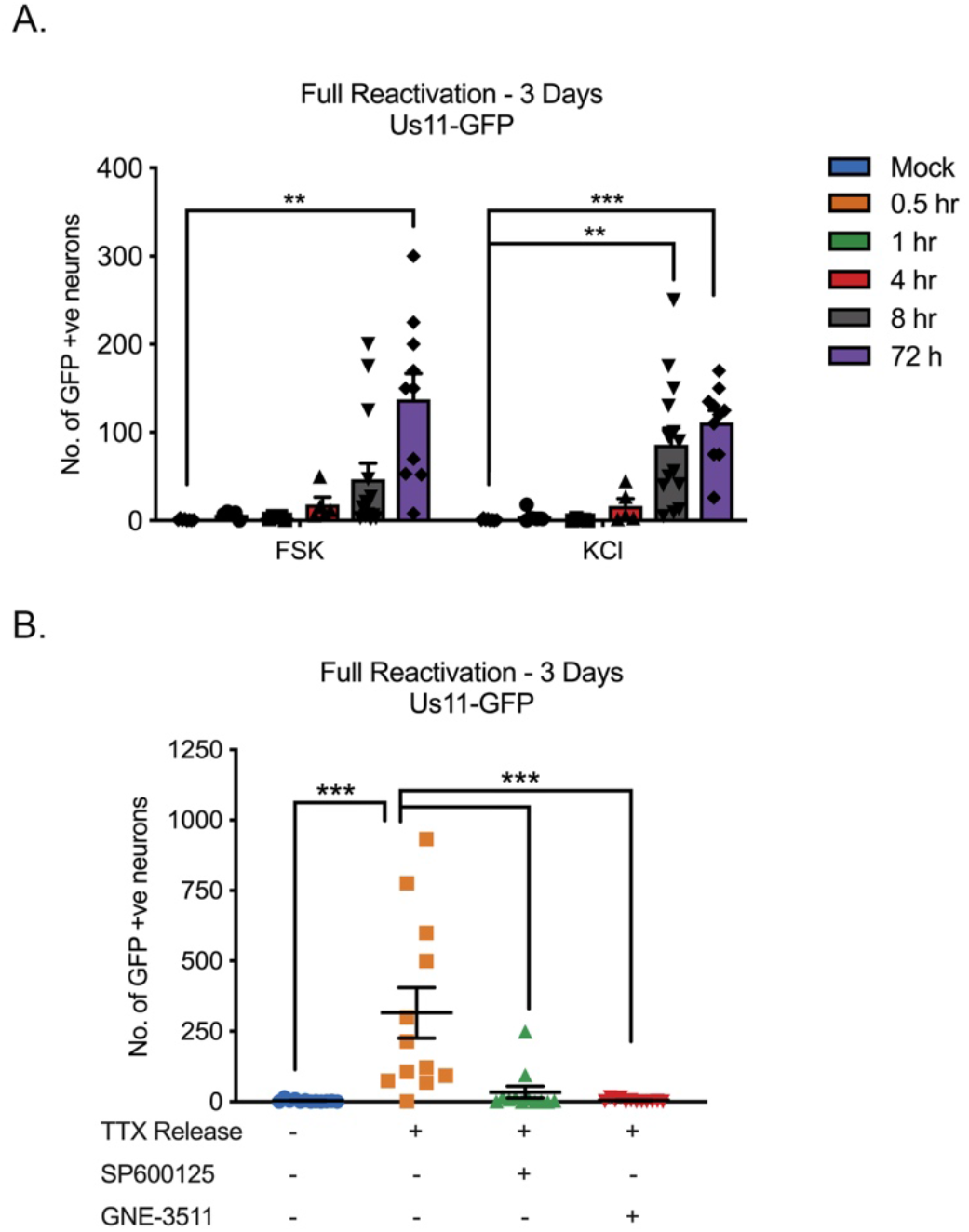
HSV Reactivation Triggered by Prolonged Neuronal Hyperexcitability is DLK/JNK Dependent. (A) Latently infected SCG cultures were treated with forskolin or KCl (55mM) for the indicated times followed by wash-out. Reactivation was quantified by number of Us11-GFP positive neurons at 3 days after the initial stimulus was added. (B) Latently infected neurons were placed in tetrodotoxin (TTX; 1μM) for 2 days and the TTX was then washed out. At the time of wash-out the JNK inhibitor SP600125 (20μM) or DLK inhibitor GNE-3511 (4μM) was added. Reactivation was quantified at 3-days post-wash-out. Individual experimental replicates are represented. Statistical comparisons were made using a one-way ANOVA with a Tukey’s multiple comparison. **P<0.01, *** P<0.001.

To further clarify that hyperexcitability can directly trigger HSV reactivation, we investigated the effects of removal from a TTX block on latently infected neurons. Addition of TTX to neurons results in synaptic scaling, so that when the TTX is removed the neurons enter a hyperexcitable state^60–63^. TTX was added to the neurons for 2 days and then washed out. This resulted in a robust HSV reactivation as determined by Us11-GFP synthesis (Figure 5B). We also investigated whether the JNK-cell stress pathway was important in HSV reactivation in response to TTX-release. Addition of the JNK inhibitor SP600125 or the DLK inhibitor GNE-3511 blocked HSV reactivation following TTX-release. Therefore, directly inducing neuronal hyperexcitability triggers HSV reactivation in a DLK/JNK-dependent manner.

### IL-1β Triggers HSV Reactivation in Mature Neurons in a DLK and HCN Channel-Dependent Manner

Our data thus far point to a reactivation of HSV following increasing episodes of neuronal hyperexcitability in a way that requires activation of the JNK-cell stress pathway. However, we wished to link this response to a physiological trigger that may stimulate HSV reactivation *in vivo*. Increased HCN-channel activity has been associated with inflammatory pain resulting from the activity of pyrogenic cytokines on neurons^64^. In addition, IL-1β is known to act on certain neurons to induce neuronal excitation^38–40^. IL-1β is released in the body during times of chronic, psychological stress. In addition, IL-1β contributes to the fever response^31–34^. In sympathetic neurons, we found that exposure of mature neurons to IL-1β induced an accumulation of the hyperexcitability-associated histone post-translational modifications γH2AX and H3K9me3/S10p (Figure 6A-C). We did not observe the same changes for post-natal neurons. The reasons for this maturation-dependent phenotype are unknown at this point but we hypothesize it could be due to changes in the expression of cellular factors required to respond to IL-1β. Therefore, these experiments were carried out on neurons that were postnatal day 28. The kinetics of induction of these histone modifications was different from what we had previously observed for forskolin treatment, as both γH2AX and H3K9me3/S10p steadily accumulated to 20h post-treatment. This likely reflects the activation of upstream signaling pathways in response to IL-1β prior to inducing neuronal excitation as IL-1β is known to increase the expression of voltage-gated sodium channels^40^. Importantly, IL-1β was able to trigger HSV reactivation in mature neurons (Figure 6D). Inhibition of voltage-gated sodium channels by TTX resulted in a significant decrease in the ability of IL-1β to induce reactivation (Figure 6E), therefore indicating that IL-1β triggered reactivation is via increasing neuronal activity. Reactivation was reduced in the presence of the HCN-channel inhibitor ZD 7288, although this decrease was not significant (P=0.2255), perhaps suggesting that IL-1β induction of neuronal activity is not directly due to the action of cAMP on HCN channels and instead HCN-channel activity may be required for maximal hyperexcitability and reactivation. Importantly, addition of the DLK inhibitor GNE-3511 blocked reactivation in response to IL-1β (Figure 6E). Therefore, IL-1β can induce HSV reactivation that is both dependent on neuronal activity and induction of the JNK neuronal cell stress response.

**Figure 6.**
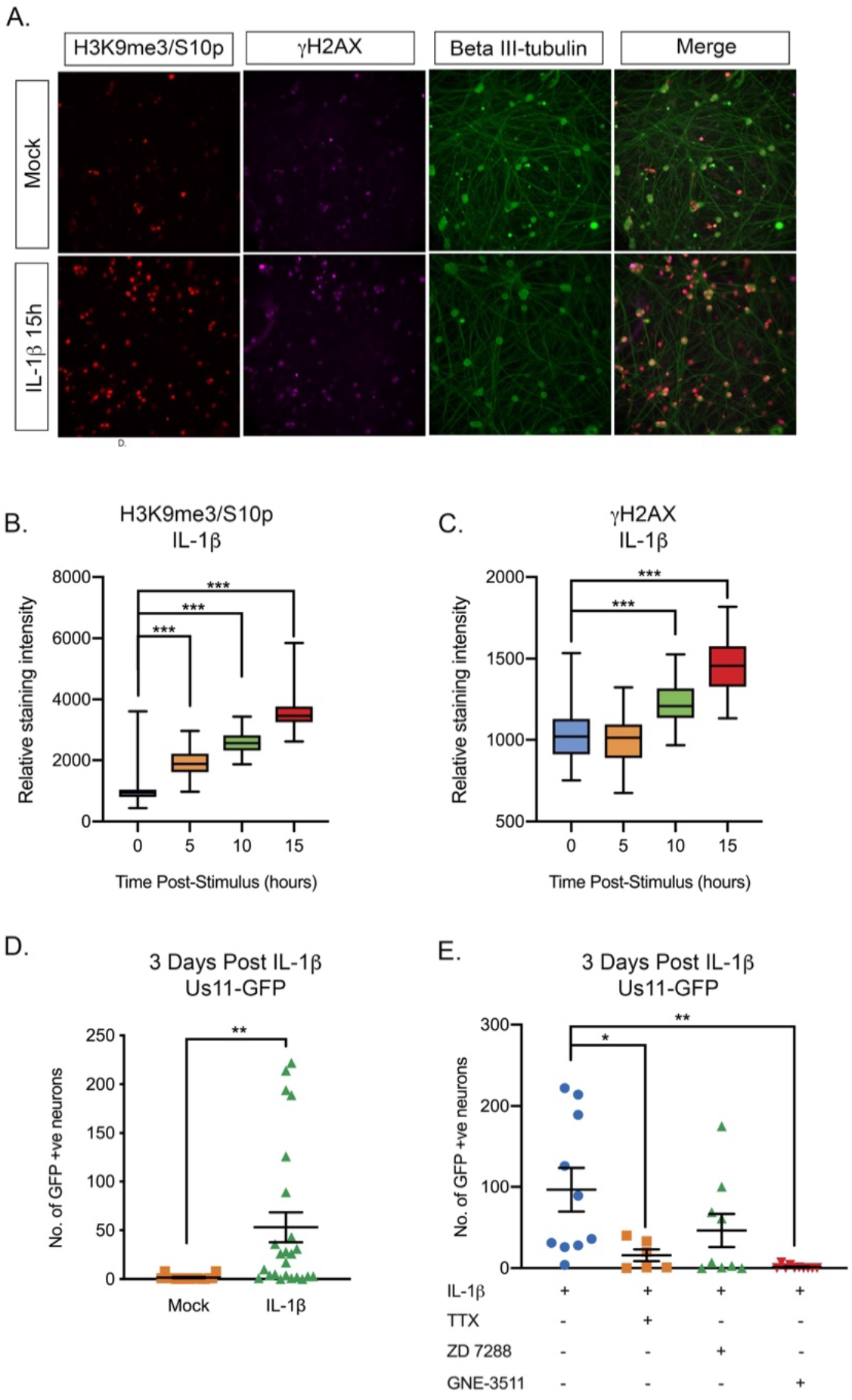
IL-1β-Induced HSV Reactivation is Linked to Heightened Neuronal Excitability and DLK Activation. (A) Adult P28 SCG neurons were treated with IL-1β (30ng/mL) for 15 hrs and stained for H3K9me3/S10p, γH2AX and beta II-tubulin to mark neurons. (B-C) Quantification of the intensity of H3K9me3/S10p and γH2AX in neuronal nuclei following forskolin treatment from two independent experiments. (D). Addition of IL-1β to latently infected cultures of mature SCG neurons triggers HSV reactivation. (E). Quantification of IL-1β induced reactivation in the presence of the voltage gated sodium channel blocker TTX (1μM), the HCN channel blocker ZD 7288 (10μM) and the DLK inhibitor GNE-3511 (4μM). In D and E individual experimental replicates are represented. Statistical comparisons were made using or two-tailed unpaired t-test (D) or a one-way ANOVA with a Tukey’s multiple comparison (B,C & E). *P<0.05, **P<0.01, *** P<0.001.

## Discussion

As herpesviruses hide in the form of a latent infection of specific cell types, they sense changes to the infected cell, resulting in the expression of viral lytic genes and ultimately reactivation. HSV establishes latency in neurons and has previously been found to respond to activation of a neuronal stress signaling pathway^15^. As an excitable cell type, the function of neurons is to rapidly transmit stimuli via the firing of action potentials, and under conditions of hyperexcitability, neurons increase their propensity to fire repeated action potentials. Here we show that this state of hyperexcitability induces HSV to undergo reactivation in a DLK/JNK dependent manner, indicating that the virus responds to both activation of cell stress signaling and prolonged hyperexcitability via a common pathway to result in reactivation. This common pathway also permits viral lytic gene expression from silenced promoters without the requirement of histone demethylase activity via a histone methyl/phospho switch. Conditions that result in hyperexcitability include prolonged periods of stress and inflammation, which are both linked to the release of IL-1β^31–34^. Consistent with this, here we show that IL-1β induces DNA damage and histone H3 phosphorylation in sympathetic neurons, which are both markers of neuronal excitability. Importantly, IL-1β triggered HSV reactivation that was dependent on neuronal activity and activation of DLK. Therefore, this study identifies a physiological stimulus that induces HSV reactivation via increasing neuronal excitability and places DLK/JNK signaling and a histone phospho/methyl switch as central to HSV reactivation.

Experiments using primary neuronal *in vitro* model systems and inducing reactivation by PI3-kinase inhibition have shown that reactivation progresses over two phases. Phase I involves the synchronous up-regulation of lytic gene expression that occurs independently of the viral transactivator VP16 and the activity of cellular histone demethylases^15,18^. A population of neurons progress to full reactivation (Phase II), which is dependent on both VP16 and HDM activity^15,18^. We previously found that lytic gene expression in Phase I is DLK/JNK dependent and is corelated with a JNK-dependent histone methyl/phospho switch on lytic gene promoters^15^. Here we demonstrate that a Phase I wave of viral gene expression that is dependent on activation of JNK but not histone demethylases also occurs in response to neuronal hyperexcitability. The co-localization of viral genomes with H3K9me3/pS10 indicates that a histone methyl/phospho switch also permits lytic gene expression to occur following forskolin treatment in a manner that is independent of HDM activity. This indicates that reactivation proceeds via a Phase I-wave of gene expression in response to multiple different stimuli. However, we note that there may be differences in the mechanism and kinetics of reactivation with different stimuli and/or strains of HSV-1 as reactivation triggered by axotomy may bypass Phase I^19,20^ and reactivation induced *in vivo* by heat shock with a more pathogenic strain of HSV triggered more rapid reactivation^65^. It will be especially interesting to determine in the future whether there are differences in the progression to reactivation with different strains of HSV. Ultimately, these reactivation kinetics may relate differences in the epigenetic structures of viral genomes that vary based on virus strains or differential manipulation of host-cell signaling pathways.

The Wilcox lab demonstrated in 1992 that reactivation can be induced by forskolin, and it has since been used as a trigger in multiple studies^25–28^. However, the mechanism by which increasing levels of cAMP induces lytic gene expression was not known. Here we link cAMP-induced reactivation to the excitation state of the neuron and show that the initial induction of viral gene expression is dependent on DLK and JNK activity but independent of CREB and PKA. The activity of PKA may be required for full reactivation, which is also consistent with a role for PKA in overcoming repression of the related Pseudorabies Virus during *de novo* axonal infection^66^. Our data also suggest that CREB may be involved in the progression to full reactivation. However, the mechanism of action of the inhibitor used here, 666-15, is not entirely clear. It has been reported as preventing CREB-mediated gene expression, but may act to prevent recruitment of histone acetyltransferases^67^. Therefore, inhibition of Phase II reactivation by 666-15 would be consistent with more large-scale chromatin remodeling on the viral genome at this stage. In addition, previous work has identified a role for inducible cAMP early repressor (ICER) in HSV reactivation^26^. ICER is a repressor of gene expression thats acts via heterodimerization with members of the CREB/ATF family of transcription factors. CREB expression is also known to be down-regulated by loss of NGF-signaling^68^, a known trigger of HSV reactivation. Therefore, it is conceivable that inhibition, rather than activation, of CREB is important for reactivation of HSV from latency.

Neuronal hyperexcitability results in DNA damage followed by repair, which together are thought to mediate the expression of cellular immediate early genes^29,30^. Here we show that forskolin treatment and IL-1β also induce DNA damage in sympathetic neurons. Previously, HSV reactivation has been found to occur following inhibition of DNA damage, inhibition of repair, and exogenous DNA damage^12^. In the context of repair inhibition or exogenous DNA damage, reactivation was dependent on dephosphorylation of AKT by the PHLPP1 phosphatase and activation of JNK, and therefore feeds into the same pathway as PI3K-inhibition. However, we did not observe decreased AKT phosphorylation in response to forskolin treatment, indicating that the mechanism of reactivation is distinct following physiological levels of DNA damage resulting from neuronal hyperexcitability versus perturbation of the damage/repair pathways.

Conditions that result in hyperexcitability include prolonged periods of stress and inflammation, which are both linked to the release of IL-1β^31–34^. Consistent with these findings, we show that IL-1β treatment induces two markers of neuronal excitability, DNA damage and histone H3 phosphorylation, in primary sympathetic neurons. The IL-1 family of cytokines act via the IL-1 receptor to activate downstream signaling pathways ^69^. IL-1β is released systemically during prolonged periods of psychological stress and upon infection via activation of the inflammasome^31–34^. IL-1α, which also signals via the IL-1R, is released locally as an alarmin. Interesting IL-1α and IL-1β are found at high levels in keratinocytes and are released upon HSV-1 infection^41^, where they can mediate antiviral responses in underlying stromal fibroblasts and endothelial cells. Antiviral responses mediated by IL-1 signaling have been found to involve NF-κB, IRF3 and/or IRF1^42^. The downstream signaling elicited by IL-1 in neurons has not been clearly defined and likely varies between different subtypes of neurons. NF-κB has been reported to be absent in certain subtypes of neurons but constitutively active in others^70,71^, and a recent study suggests that NF-κB levels increase with neuronal maturation^72^, which may be why we only observed IL-1-mediated reactivation in mature neurons. Additional studies have found a role for p38MAPK signaling and AKT/mTOR signaling in neuronal IL-1-mediated responses^73,74^. A common feature of IL-1 signaling in neurons is increased excitability, which has been associated with neurotransmitter release, and mediates a variety of physiological responses including behavior modulation and an intersection with the hosts’ immune response^38–40^. IL-1 is also associated with pathological conditions, including neurodegenerative disease such as Alzheimer’s disease^75^. There are multiple studies linking HSV-1 infection to the progression of Alzheimer’s^2^; therefore, the combination of both HSV infection and increased IL-1 could have a feed forward effect on the progression of Alzheimer’s disease by promoting increased reactivation of HSV from latency.

Previously, we found that JNK activation by DLK is required for reactivation following interruption of the NGF-signaling pathway. Here we find that forskolin and IL-1β-mediated reactivation also required both DLK activity, further reinforcing the central role of DLK and JNK in reactivation of HSV from latency. DLK is known as a master regulator of neuronal response to stress stimuli and mediates whole cell death, axon pruning, regeneration or generation depending on the nature of the stimuli. However, it has not before been linked to neuronal hyperexcitability or the response to IL-1β signaling. The known mechanisms of DLK activation include loss of AKT activation and phosphorylation by PKA^52,76^, neither of which could be linked to HSV reactivation mediated by forskolin in this study. Following activation by DLK, one mechanism by which JNK is thought to permit lytic gene expression is via recruitment to viral promoters and histone phosphorylation. However, it is likely that there are additional, JNK-dependent effects including activation of pioneer or transcription factors that also mediate viral gene expression. Further insight into how HSV has hijacked this cellular pathway to induce lytic gene expression may lead to novel therapeutics that prevent reactivation, in addition to providing information on how viral gene expression initiates from promoters assembled into heterochromatin.

---

**Figure S1.**
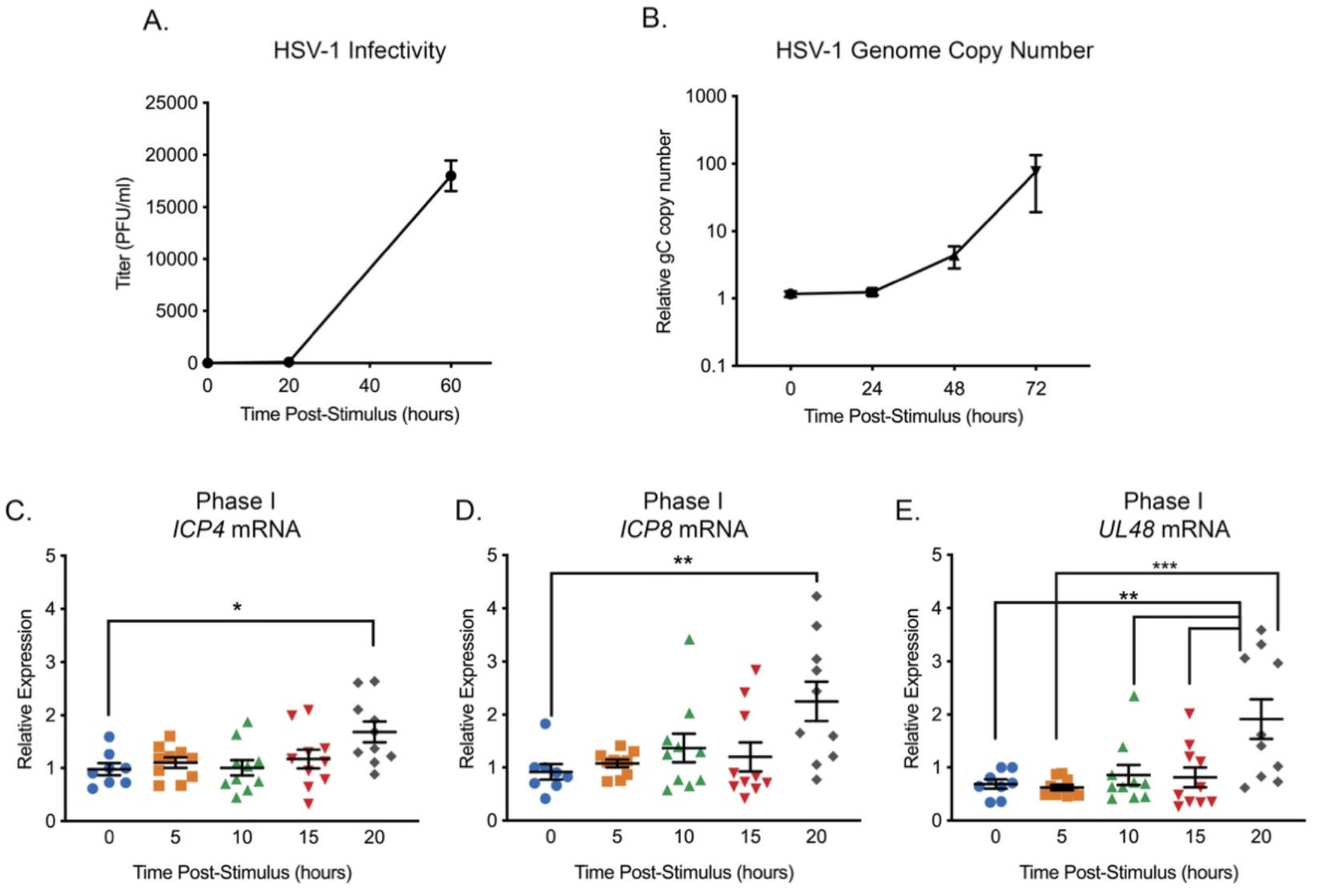
Reactivation Triggered by Forskolin Triggers a Wave of Lytic Gene Expression that Precedes DNA Replication and Infectious Virus Production (supplement to Figure 2) (A) Titers of infectious virus detected from reactivating neurons induced with forskolin (n=4). (B) Quantification of the relative viral genome copy number following forskolin-mediated reactivation based on *gC* copy number normalized to cellular GAPDH and expressed relative to the 0h time-point (n=7). (C-E) RT-qPCR for viral mRNA transcripts following forskolin treatment of latently infected SCGs. In C-E each biological replicate is represented. Statistical comparisons were made using a one-way ANOVA with a Tukey’s multiple comparison. *P<0.05, ** P<0.01, ***P<0.001. (C-E).

**Figure S2.**
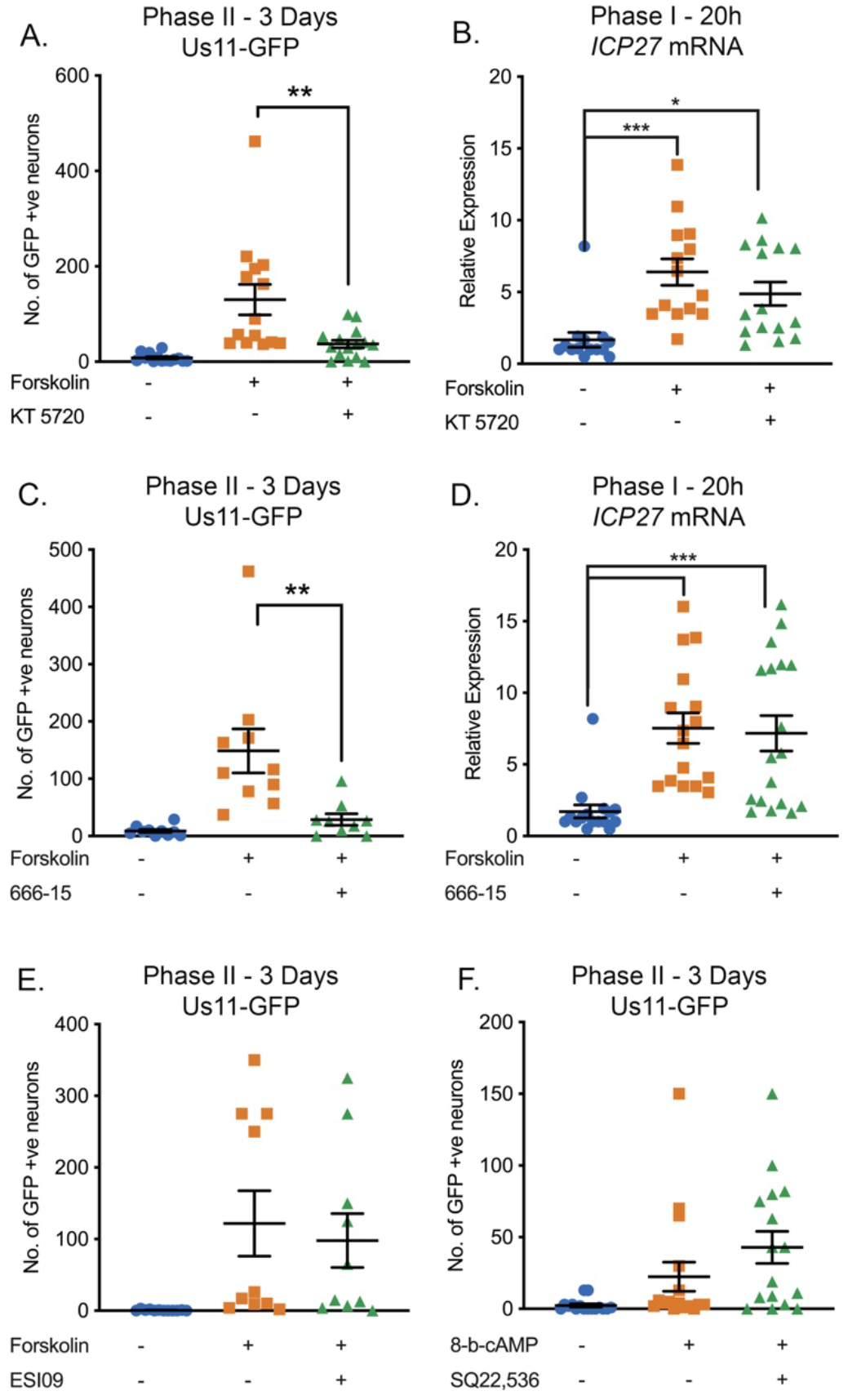
Effect of PKA, CREB, Rapgef2 and EPAC Inhibition on HSV-1 Reactivation. (A) Latently infected cultures were reactivated with forskolin (60 μM) in the presence of the PKA inhibitor KT 5720 (3 μM) and the number of Us11-GFP positive neurons quantified at 3 days post-reactivation. (B) RT-qPCR for the viral lytic transcript ICP27 at 20h post-forskolin treatment and in the presence of KT 5720. (C) Latently infected cultures were reactivated with forskolin in the presence of the CREB inhibitor 666-15 (2 μM). (D) RT-qPCR for ICP27 at 20h post-forskolin treatment and in the presence of 666-15. (E) Latently infected cultures were reactivated with forskolin (60μM) in the presence of the EPAC inhibitor ESI09 (10 μM). (F) Latently infected cultures were reactivated with 8-Bromo-cAMP (125μM) in the presence of the Rapgef2 inhibitor SQ22,536 (50 μM). Individual experimental replicates are represented. Statistical comparisons were made using a one-way ANOVA with a Tukey’s multiple comparison. *P<0.05, ** P<0.01, ***P<0.001.

**Figure S3.**
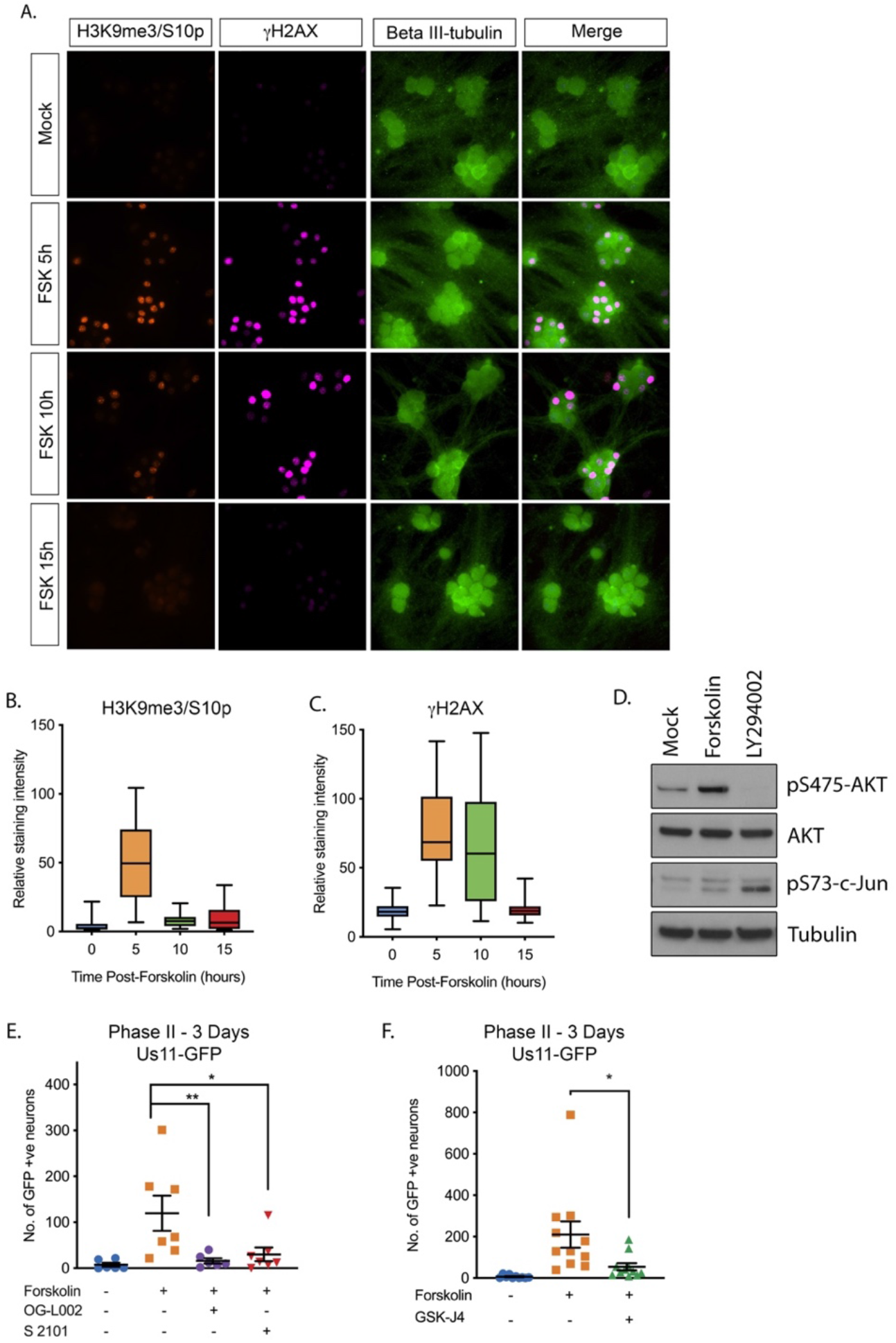
Forskolin Treatment Induces Hyperexcitability-associated Histone Post-translational Modification in Sympathetic Neurons. SCG neurons were treated with forskolin and immunofluorescence staining was carried out for H3K9me3/S10p, the DNA damage marker γH2AX and the neuronal marker beta III-tubulin. (B) Quantification of neuronal nuclear staining intensity for H3K9me3 (>150 cells/condition). (C) Quantification of neuronal nuclear staining for γH2AX. In B and C, whiskers represent the 2.5-97.5 percentile range. (D). Western blotting for pS475-AKT, total AKT, pS73-c-Jun and tubulin at 15h post-treatment with the PI3-kinase inhibitor LY294002 (20μM) or forskolin (60 μM). (E) Effect of the LSD1 inhibitors OG-L002 (15μM) and S 2101 (20μM) on forskolin-mediated reactivation measured by Us11-GFP positive neurons. (F) Effect of the JMJD3 and UTX inhibitor GSK-J4 (2μM) on forskolin-mediated reactivation measured by Us11-GFP positive neurons. Statistical comparisons were made using a one-way ANOVA with a Tukey’s multiple comparison. *P<0.05, ** P<0.01, ***P<0.001 (E, F).

**Figure S4.**
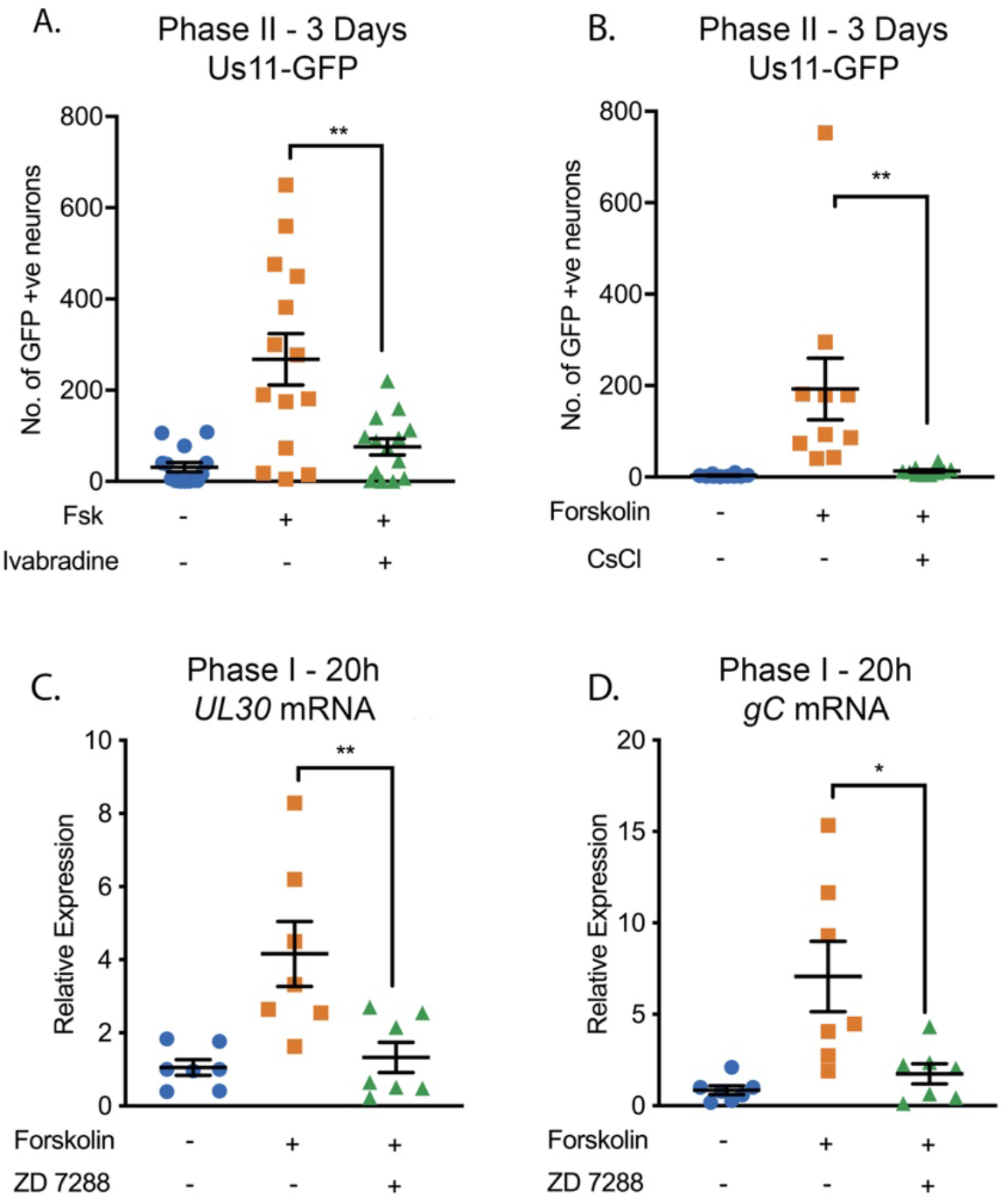
HSV Reactivation Mediated by Forskolin Requires Neuronal Excitability. (A and B) Latently infected cultures were reactivated with forskolin in the presence of the HCN channel inhibitors ivabradine (20μM; A) and CsCl (3mM; B). Latently infected cultures were reactivated with forskolin in the presence of the HCN inhibitor ZD 7288 (10 μM) and viral lytic transcripts measured at 20h post-reactivation (C and D). Individual experimental replicates are represented. Statistical comparisons were made using a one-way ANOVA with a Tukey’s multiple comparison. *P<0.05, ** P<0.01.

## Acknowledgements

We thank Ian Mohr (NYU) for supplying the Us11-GFP virus used in this study. This work was supported by NIH/NINDS R01NS105630 (to A.R.C), NIH/NIAID T32AI007046 (S.R.C. and J.B.S), NIH/NEI F30EY030397 (J.B.S), NIH/NIGMS T32GM008136 (S.D) and T32GM007267 (J.B.S) and MRC (https://mrc.ukri.org) MC_UU_12014/5 (C.B).

## Materials and Methods

### Reagents

Compounds used in the study are as follows: Acycloguanosine, FUDR, Uridine, SP600125, GNE-3511, GSK-J4, L-Glutamic Acid, and Ivabradine (Millipore Sigma); Forskolin, LY 294002, 666-15, SQ 22536, KT 5720, Tetraethylammonium chloride, Cesium chloride, OG-L002, S2101, Tetrotdotoxin, and ESI-09 (Tocris); 1,9-dideoxy Forskolin, ZD 7288 and 8-bromo-cyclic AMP (Cayman Chemicals); Nerve Growth Factor 2.5S (Alomone Labs); Primocin (Invivogen); Aphidicolin (AG Scientific); IL-1β (Shenandoah Biotechnology); WAY-150138 was kindly provided by Pfizer, Dr. Jay Brown and Dr. Dan Engel at the University of Virginia, and Dr. Lynn Enquist at Princeton University. Compound information and concentrations used can be found below in Table S1. Compound concentrations were used based on previously published IC50s and assessed for neuronal toxicity using the cell body and axon health and degeneration index (Table S5 and S6). All compounds used had an average score ≤1.

### Preparation of HSV-1 Virus Stocks

HSV-1 stocks of eGFP-Us11 Patton were grown and titrated on Vero cells obtained from the American Type Culture Collection (Manassas, VA). Cells were maintained in Dulbecco’s Modified Eagle’s Medium (Gibco) supplemented with 10% FetalPlex (Gemini Bio-Products) and 2 mM L-Glutamine. eGFP-Us11 Patton (HSV-1 Patton strain with eGFP reporter protein fused to true late protein Us11^43^) was kindly provided by Dr. Ian Mohr at New York University.

### Primary Neuronal Cultures

Sympathetic neurons from the Superior Cervical Ganglia (SCG) of post-natal day 0-2 (P0-P2) or adult (P21-P24) CD1 Mice (Charles River Laboratories) were dissected as previously described^15^. Rodent handling and husbandry were carried out under animal protocols approved by the Animal Care and Use Committee of the University of Virginia (UVA). Ganglia were briefly kept in Leibovitz’s L-15 media with 2.05 mM L-Glutamine before dissociation in Collagenase Type IV (1 mg/mL) followed by Trypsin (2.5 mg/mL) for 20 minutes each at 37 °C. Dissociated ganglia were triturated, and approximately 10,000 neurons per well were plated onto rat tail collagen in a 24-well plate. Sympathetic neurons were maintained in CM1 (Neurobasal® Medium supplemented with PRIME-XV IS21 Neuronal Supplement (Irvine Scientific), 50 ng/mL Mouse NGF 2.5S, 2 mM L-Glutamine, and Primocin). Aphidicolin (3.3 μg/mL), Fluorodeoxyuridine (20 μM) and Uridine (20 μM) were added to the CM1 for the first five days post-dissection to select against proliferating cells.

### Establishment and Reactivation of Latent HSV-1 Infection in Primary Neurons

Latent HSV-1 infection was established in P6-8 sympathetic neurons from SCGs. Neurons were cultured for at least 24 hours without antimitotic agents prior to infection. The cultures were infected with eGFP-Us11 (Patton recombinant strain of HSV-1 expressing an eGFP reporter fused to true late protein Us11). Neurons were infected at a Multiplicity of Infection (MOI) of 7.5 PFU/cell (assuming 1.0×10^4^ neurons/well/24-well plate) in DPBS +CaCl2 +MgCl2 supplemented with 1% Fetal Bovine Serum, 4.5 g/L glucose, and 10 μM Acyclovir (ACV) for three hours at 37 °C. Post-infection, inoculum was replaced with CM1 containing 50 μM ACV for 5-6 days, followed by CM1 without ACV. Reactivation was carried out in DMEM/F12 (Gibco) supplemented with 10% Fetal Bovine Serum, Mouse NGF 2.5S (50 ng/mL) and Primocin. Inhibitors were added either one hour prior to or concurrently with the reactivation stimulus. WAY-150138 (2-10 μg/mL) was added to reactivation cocktail to limit cell-to-cell spread. Reactivation was quantified by counting number of GFP-positive neurons or performing Reverse Transcription Quantitative PCR (RT-qPCR) of HSV-1 lytic mRNAs isolated from the cells in culture.

### Analysis of mRNA expression by reverse-transcription quantitative PCR (RT-qPCR)

To assess relative expression of HSV-1 lytic mRNA, total RNA was extracted from approximately 1.0×10^4^ neurons using the Quick-RNA™ Miniprep Kit (Zymo Research) with an on-column DNase I digestion. mRNA was converted to cDNA using the SuperScript IV First-Strand Synthesis system (Invitrogen) using equal amounts of RNA (20-30 ng/reaction). To assess viral DNA load, total DNA was extracted from approximately 1.0×10^4^ neurons using the Quick-DNA™ Miniprep Plus Kit (Zymo Research). qPCR was carried out using *Power* SYBR™ Green PCR Master Mix (Applied Biosystems). The relative mRNA or DNA copy number was determined using the Comparative C_T_ (ΔΔC_T_) method normalized to mRNA or DNA levels in latently infected samples. Viral RNAs were normalized to mouse reference gene GAPDH. All samples were run in duplicate on an Applied Biosystems™ QuantStudio™ 6 Flex Real-Time PCR System and the mean fold change compared to the reference gene calculated. Primers used are described in Table S2.

### Western Blot Analysis

Neurons were lysed in RIPA Buffer with cOmplete, Mini, EDTA-Free Protease Inhibitor Cocktail (Roche) and PhosSTOP Phosphatase Inhibitor Cocktail (Roche) on ice for one hour with regular vortexing to aid lysis. Insoluble proteins were removed via centrifugation, and lysate protein concentration was determined using the Pierce Bicinchoninic Acid Protein Assay Kit (Invitrogen) using a standard curve created with BSA standards of known concentration. Equal quantities of protein (generally 20-50 μg) were resolved on 4-20% gradient SDS-Polyacrylamide gels (Bio-Rad) and then transferred onto Polyvinylidene difluoride membranes (Millipore Sigma). Membranes were blocked in PVDF Blocking Reagent for Can Get Signal (Toyobo) for one hour. Primary antibodies were diluted in Can Get Signal Immunoreaction Enhancer Solution 1 (Toyobo) and membranes were incubated overnight at 4°C. Antibodies and concentrations are described in Table S3 below. HRP-labeled secondary antibodies were diluted in Can Get Signal Immunoreaction Enhancer Solution 2 (Toyobo) and membranes were incubated for one hour at room temperature. Blots were developed using Western Lightning Plus-ECL Enhanced Chemiluminescence Substrate (PerkinElmer) and ProSignal ECL Blotting Film (Prometheus Protein Biology Products) according to manufacturer’s instructions. Blots were stripped for reblotting using NewBlot PVDF Stripping Buffer (Licor).

### Immunofluorescence

Neurons were fixed for 15 minutes in 4% Formaldehyde and blocked in 5% Bovine Serum Albumin and 0.3% Triton X-100 and incubated overnight in primary antibody. Antibodies and concentrations are described in Table S4 below. Following primary antibody treatment, neurons were incubated for one hour in Alexa Fluor® 488-, 555-, and 647-conjugated secondary antibodies for multi-color imaging (Invitrogen). Nuclei were stained with Hoechst 33258 (Life Technologies). Images were acquired using an sCMOS charge-coupled device camera (pco.edge) mounted on a Nikon Eclipse Ti Inverted Epifluorescent microscope using NIS-Elements software (Nikon). Images were analyzed and intensity quantified using ImageJ.

### Click Chemistry

Click chemistry was carried out a described previously^53^ with some modifications. Neurons were washed with CSK buffer (10 mM HEPES, 100 mM NaCl, 300 mM Sucrose, 3 mM MgCl2, 5 mM EGTA) and simultaneously fixed and permeabilized for 10 minutes in 1.8% methonal-free formaldehyde (0.5% Triton X-100, 1% phenylmethylsulfonyl fluoride (PMSF)) in CSK buffer, then washed twice with PBS before continuing to the click chemistry reaction and immunostaining. Samples were blocked with 3% BSA for 30 minutes, followed by click chemistry using EdC-labelled HSV-1 DNA and the Click-iT EdU Alexa Flour 555 Imaging Kit (ThermoFisher Scientific, C10638) according to the manufacturer’s instructions. For immunostaining, samples were incubated overnight with primary antibodies in 3% BSA. Following primary antibody treatment, neurons were incubated for one hour in Alexa Fluor® 488-, 555-, and 647-conjugated secondary antibodies for multi-color imaging (Invitrogen). Nuclei were stained with Hoechst 33258 (Life Technologies). Images were acquired at 60x using an sCMOS charge-coupled device camera (pco.edge) mounted on a Nikon Eclipse Ti Inverted Epifluorescent microscope using NIS-Elements software (Nikon). Images were analyzed and intensity quantified using ImageJ.

### Statistical Analysis

Power analysis was used to determine the appropriate sample sizes for statistical analysis. All statistical analysis was performed using Prism V8.4. An unpaired t-test was used for all experiments where the group size was 2. All other experiments were analyzed using a one-way ANOVA with a Tukey’s multiple comparison. Specific analyses are included in the figure legends. For all reactivation experiments measuring GFP expression, viral DNA, gene expression or DNA load, individual biological replicates were plotted (an individual well of primary neurons) and all experiments were repeated from pools of neurons from at least 3 litters. EdC virus and H3K9me3S10/p co-localization was quantified using ImageJ after sample blinding of at least 8 fields of view from 2 biological replicates. Mean fluorescence intensity of γH2AX and H3K9me3pS10 was quantified using ImageJ from at least 100 cells from at least 3 biological replicates.

## Supplemental Materials and Methods Tables

**Table S1:** Compounds Used and Concentrations

| Compound | Supplier | Identifier | Concentration |
| --- | --- | --- | --- |
| Acycloguanosine | Millipore Sigma | A4669 | 10 μM, 50 μM |
| FUDR | Millipore Sigma | F-0503 | 20 μM |
| Uridine | Millipore Sigma | U-3003 | 20 μM |
| SP600125 | Millipore Sigma | S5567 | 20 μM |
| GNE-3511 | Millipore Sigma | 533168 | 4 μM |
| GSK-J4 | Millipore Sigma | SML0701 | 2 μM |
| L-Glutamic Acid | Millipore Sigma | G5638 | 3.7 μg/mL |
| Forskolin | Tocris | 1099 | 60 μM |
| LY 294002 | Tocris | 1130 | 20 μM |
| 666-15 | Tocris | 5661 | 2 μM |
| SQ 22,536 | Tocris | 1435 | 50 μM |
| KT 5720 | Tocris | 1288 | 3 μM |
| TEA | Tocris | 3068 | 10 mM |
| CsCl | Tocris | 4739 | 3 mM |
| OG-L002 | Tocris | 6244 | 15 μM, 30 μM |
| S2101 | Tocris | 5714 | 10 μM, 20 μM |
| Tetrodotoxin | Tocris | 1069 | 1 μM |
| ESI-09 | Tocris | 4773 | 10 μM |
| ZD 7288 | Cayman | 15228 | 20 μM |
| 8-bromo-cyclic AMP | Cayman | 14431 | 125 µM |
| NGF 2.5S | Alomone Labs | N-100 | 50 ng/mL |
| Primocin | Invivogen | ant-pm-1 | 100 µg/mL |
| Aphidicolin | AG Scientific | A-1026 | 3.3 µg/mL |
| IL-1β | Shenandoah Bio. | 100-167 | 30ng/mL |
| WAY-150138 | Pfizer | NA | 10 µg/mL |

**Table S2:**
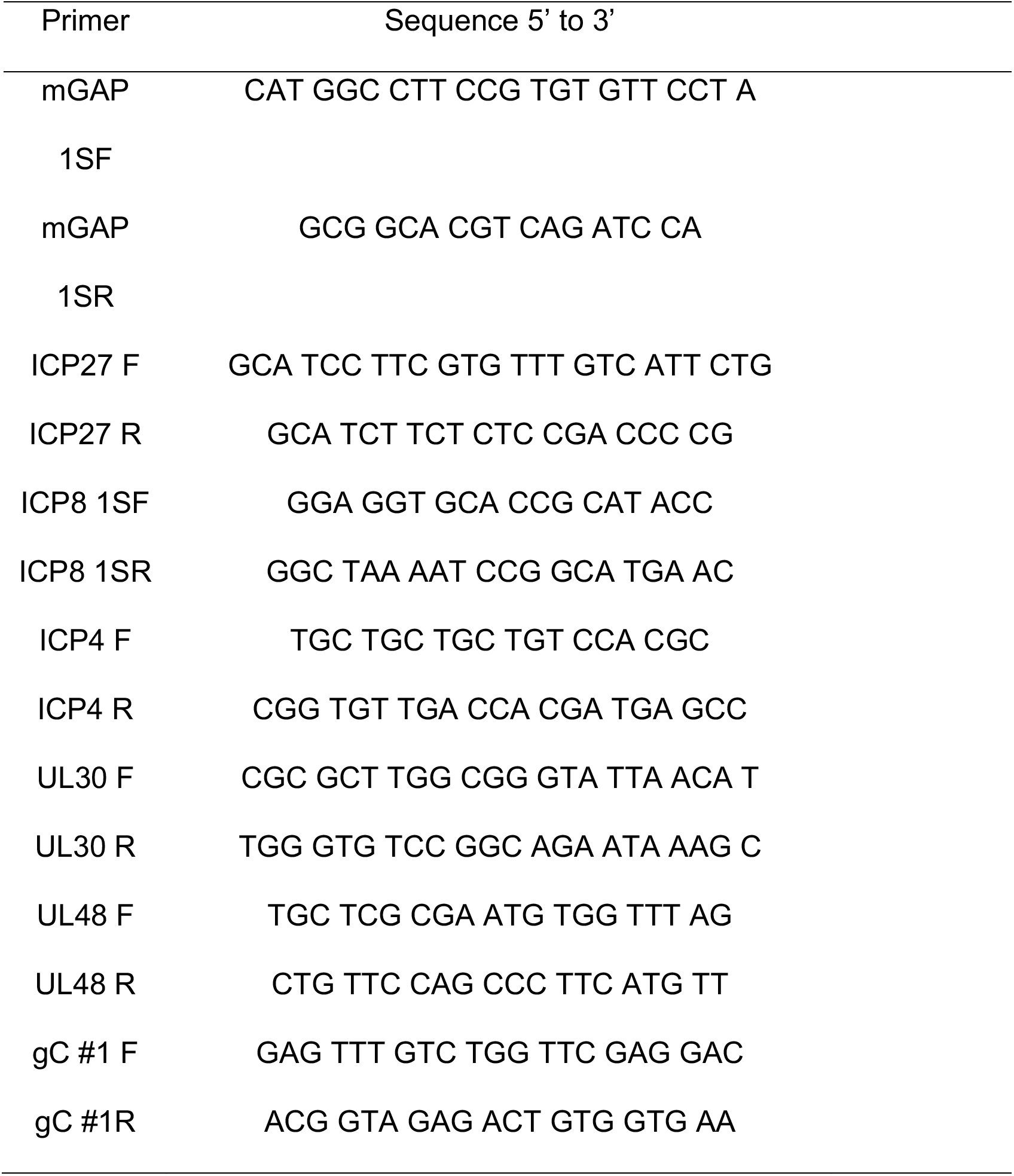
Primers Used for RT-qPCR

**Table S3:**
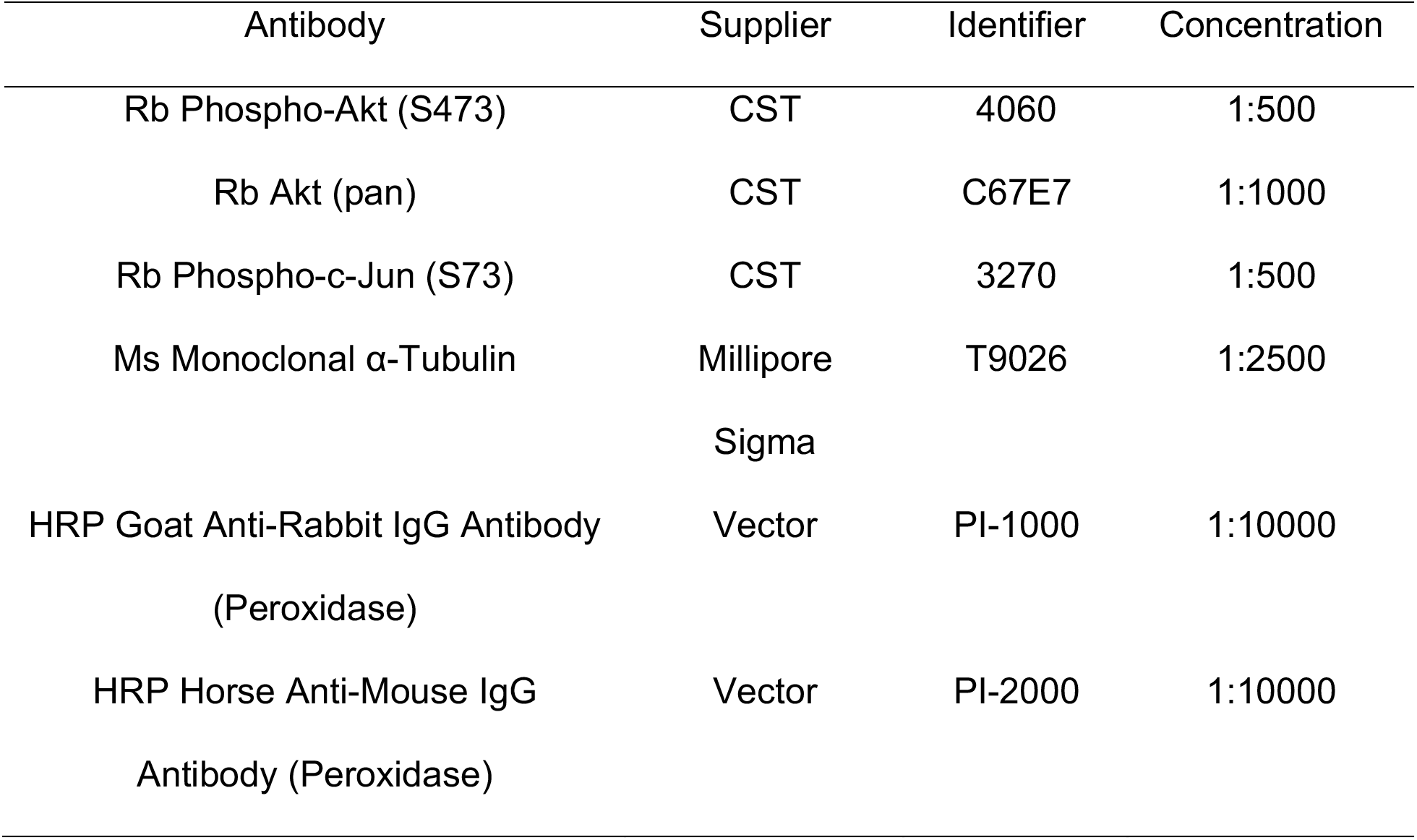
Antibodies Used for Western Blotting and Concentrations

**Table S4:** Antibodies Used for Immunofluorescence and Concentrations

| Antibody | Supplier | Identifier | Concentration |
| --- | --- | --- | --- |
| Rb H3K9me3S10P | Abcam | ab5819 | 1:250 |
| Ch Beta-III Tubulin | Millipore sigma | AB9354 | 1:1000 |
| Ms γH2A.X | CST | 80312S | 1:100 |
| Ms c-Fos | Novus | NB110-75039 | 1:125 |
| F(ab')<sub>2</sub> Goat anti Mouse IgG (H+L) Alexa Fluor® 647 | Thermo Fisher | A21237 | 1:1000 |
| F(ab') Goat anti Rabbit IgG (H+L) Alexa Fluor® 555 | Thermo Fisher | A21425 | 1:1000 |
| Goat anti Chicken IgY (H+L) Alexa Fluor® 647 | abcam | ab150175 | 1:1000 |
| Goat Anti-Chicken IgY H&L (Alexa Fluor® 488) preabsorbed | abcam | ab150173 | 1:1000 |
| F(ab')<sub>2</sub> Goat anti-Rabbit IgG (H+L) Alexa Fluor® 488) | Thermo Fisher | B40922 | 1:1000 |

**Table S5:** Cell Body Score for Neuronal Health and Degeneration Index

| Score | Description |
| --- | --- |
| 0 | Large, phase bright cell bodies. Clear with no fragmentation or vesiculation. |
| 1 | Small, phase bright cell bodies. Clear with no fragmentation or vesiculation. |
| 2 | Cell bodies do not have fragmentation but are not phase bright. Sometimes appear transparent. |
| 3 | Cell bodies with fragmentation but few dead neurons or corpses. |
| 4 | Cell bodies with fragmentation with many corpses present and neurons starting to detach. |
| 5 | Complete cell death. Neurons detached. |

**Table S6:** Axon Score for Neuronal Health and Degeneration Index

| Score | Description |
| --- | --- |
| 0 | Axons totally smooth with no blebbing or fragmentation. Branched and form a spider web-like network. |
| 1 | Axons smooth but grow straight. |
| 2 | Blebbing on the axons but no apparent fragmentation. |
| 3 | Fragmentation starting to appear in <50% of the neurons. |
| 4 | Fragmentation in >50% of the neurons. |
| 5 | No axons remaining. |

